# SVJedi: Genotyping structural variations with long reads

**DOI:** 10.1101/849208

**Authors:** Lolita Lecompte, Pierre Peterlongo, Dominique Lavenier, Claire Lemaitre

## Abstract

**Motivation:** Studies on structural variants (SV) are expanding rapidly. As a result, and thanks to third generation sequencing technologies, the number of discovered SVs is increasing, especially in the human genome. At the same time, for several applications such as clinical diagnoses, it is important to genotype newly sequenced individuals on well defined and characterized SVs. Whereas several SV genotypers have been developed for short read data, there is a lack of such dedicated tool to assess whether known SVs are present or not in a new long read sequenced sample, such as the one produced by Pacific Biosciences or Oxford Nanopore Technologies.

**Results:** We present a novel method to genotype known SVs from long read sequencing data. The method is based on the generation of a set of reference sequences that represent the two alleles of each structural variant. Long reads are aligned to these reference sequences. Alignments are then analyzed and filtered out to keep only informative ones, to quantify and estimate the presence of each SV allele and the allele frequencies. We provide an implementation of the method, SVJedi, to genotype insertions and deletions with long reads. The tool has been applied to both simulated and real human datasets and achieves high genotyping accuracy. We also demonstrate that SV genotyping is considerably improved with SVJedi compared to other approaches, namely SV discovery and short read SV genotyping approaches.

**Availability:** https://github.com/llecompte/SVJedi.git

## 1 Introduction

Structural variations (SVs) are characterized as genomic segments of at least 50 base pair (bp) long, that are rearranged in the genome. There are several types of SV such as deletions, insertions, duplications, inversions or translocations. With the advent of Next-Generation Sequencing (NGS) technologies and the re-sequencing of many individuals in populations, SVs have been admitted as a key component of human polymorphism [1]. This kind of polymorphism has been shown involved in many biological processes such as diseases or evolution [2]. Databases referencing such variants grow as new variants are discovered. At this time, dbVar, the reference database of human genomic SVs [3] now contains more than 36 million variant calls, illustrating that many SVs have already been discovered and characterized in human populations.

When studying SVs in newly sequenced individuals, one can distinguish two distinct problems: discovery and genotyping. In the SV discovery problem, the aim is to identify all the variants that differentiate the given re-sequenced individual with respect usually to a reference genome. In the SV genotyping problem, the aim is to evaluate if a given known SV (or set of SVs) is present or absent in the re-sequenced individual, and assess, if it is present, with which ploidy (heterozygous or homozygous). At first glance, the genotyping problem may seem included in the discovery problem, since present SVs should be discovered by discovery methods. However, in discovery algorithms, SV evidence is only investigated for present variants (*i.e.* incorrect mappings) and not for absent ones. If a SV has not been called, we cannot know if the caller missed it (false negative) or if the variant is truly absent in this individual and this could be validated by a significant amount of correctly mapped reads in this region. Moreover, in the genotyping problem, knowing what we are looking for, should make the problem simpler and the genotyping result hopefully more precise. With the fine characterization of a growing number of SVs in populations of many organisms, genotyping newly sequenced individuals becomes very interesting and informative, in particular in human medical diagnosis contexts or more generally in any association or population genomics studies.

In this work, we focus on the second problem: genotyping already known SVs in a newly sequenced sample. Such genotyping methods already exist for short reads data: for instance, SVtyper [4], SV^2^ [5], Nebula [6]. Though short reads are often used to discover and genotype SVs, this is well known that their short size makes them ill-adapted for predicting large SVs or SVs located in repeated regions. SVs are often located alongside repeated sequences such as mobile elements, resulting in mappability issues that make the genotyping problem harder when using short read data [7].

Third generation sequencing technology, such as Pacific Biosciences (PacBio) and Oxford Nanopore Technologies, can produce much longer reads compared to NGS technologies. Despite their high error rate, long reads are crucial in the study of SVs and have enabled new SV discoveries [8, 9, 10, 11]. Indeed, the size range of these sequences can reach a few kilobases (kb) to megabases, thus long reads can extend over rearranged sequence portions as well as over the repeated sequences often present at SV’s breakpoint regions.

Following long read technology’s development, many SV discovery tools have emerged, such as Sniffles [12], NanoSV [10] or SVIM [13]. Among these tools, some implement a genotyping module that gives the frequency of alleles after calling SVs of the sequenced samples. Nonetheless, they require post-processing to evaluate if a set of SVs is present or not in the sample and to compare the SV calls between different samples. To our knowledge, there is currently no dedicated tool that can perform SV genotyping with long read data.

In this work, we focused on the vast majority and most studied types of SVs: the deletions and insertions. These two types of variants represent 99 % of SVs characterized in the human genomic SV database, dbVar (including duplications and mobile element movements as sub-types of insertions and deletions). In particular, in the clinical diagnosis context, such unbalanced variants and especially deletions are primarily investigated compared to other SV types because of their clear functional impact.

The main contribution of this work is a novel method to genotype known deletions and insertions using long read data. We also provide an implementation of this method in the tool named SVJedi. SVJedi accuracy and robustness were evaluated on simulated data of real deletions in a human chromosome. It was also applied to a real human dataset and compared to a gold standard call set provided by the Genome in a Bottle (GiaB) Consortium, containing both deletions and insertions. High precision was achieved on both simulated and real data. We also demonstrated the improvement of such a dedicated method over other approaches, namely SV discovery with long reads and SV genotyping with short reads.

## 2 Methods

The method assigns a genotype for a set of already known SVs in a given individual sample sequenced with long read data. It assesses for each SV if it is present in the given individual, and if so, how many variant alleles it holds, *i.e.* whether the individual is heterozygous or homozygous for the particular variant.

For clarity purposes, we describe here the method for deletion genotyping only. The genotyping of insertions, also implemented in SVJedi, is perfectly symmetrical. The method takes as input a variant file with deletion coordinates in VCF format, a reference genome and the sample of long read sequences. It outputs a variant file complemented with the individual genotype information for each input variant in VCF format.

The method consists of four different steps, that are illustrated in Figure 1. The fundamentals of the method lie in its first step, which generates reference sequences that represent the two alleles of each SV. Long reads are then aligned on the whole set of reference alleles. An important step consists of selecting and counting only informative alignments to finally estimate the genotype for each input variant.

**Figure 1:**
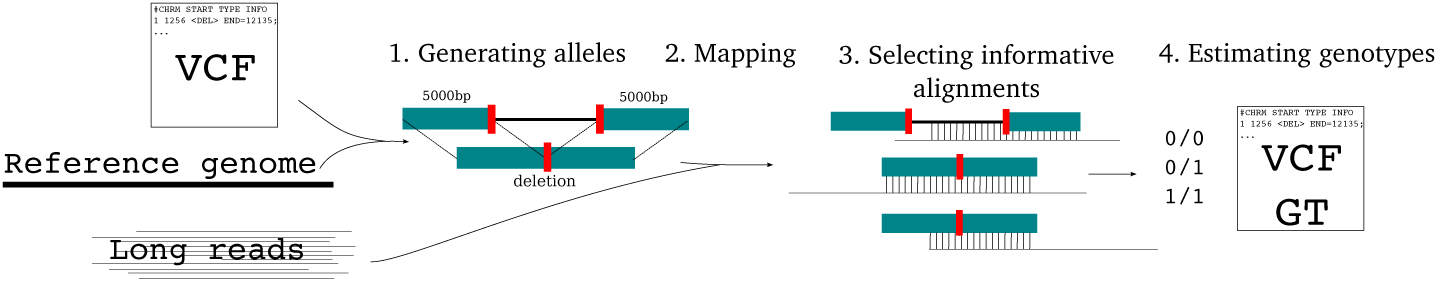
SVJedi steps for deletion genotyping. Steps for insertion genotyping are symmetrical and are not shown on the figure for clarity purposes. 1. Two corresponding reference sequences are generated for each selected SV, one corresponds to the original sequence and the other to the sequence with the deletion. 2. Long read sequenced data are aligned on these references using Minimap2. 3. Informative alignments are selected. 4. Genotypes are estimated.

### 2.1 Allele sequences generation

Starting from a known variant file in VCF format and the corresponding reference genome, the first step consists of generating two sequences for each SV, corresponding to the two possible allele sequences. In the case of deletions, these are parts of the reference genome that may be absent in a given individual. They are characterized in the VCF file by a starting position on the reference genome and a length. We define the reference allele sequence (allele 0) as the sequence of the deletion with adjacent sequences at each side, and the alternative allele (allele 1) consists of the joining of the two previous adjacent sequences. Given that reads of several kb will be mapped on these reference sequences, the size of the adjacent sequences was set to 5,000 bp at each side, giving a 10 kb sequence for allele 1 and 10 kb plus the deletion size for allele 0. For deletions larger than 10 kb, two sequence references are generated for allele 0, one for each breakpoint. The same adjacent sequence size is used, *i.e.* 5,000 bp, on each side of the breakpoints, giving then three 10 kb sequences: one for allele 1, and two for allele 0 (left and right breakpoints).

### 2.2 Mapping

Sequenced long reads are aligned on all previously generated references, using Minimap2 [14] (version 2.17-r941). Option `-c` is specified to generate a CIGAR for each alignment. Alignments are then output in a PAF file.

### 2.3 Informative alignment selection

Minimap2 raw alignment results have to be carefully filtered out to remove i) uninformative alignments, which are those not discriminating between the two possible alleles, and ii) spurious false positive alignments, that are mainly due to repeated sequences.

Informative alignments for the genotyping problem are those that overlap the SV breakpoints, that is the sequence adjacencies that are specific to one or the other allele. In the case of a deletion, the reference allele contains two such breakpoints, the start and end positions of the deletion sequence; the alternative sequence, the shortest one, contains one such breakpoint at the junction of the two adjacent sequences (see the red thick marks of Figure 1).

To be considered as overlapping a breakpoint, an alignment must cover at least *dover* bp from each side of the breakpoint (*dover* is set by default to 100 bp). In other words, if *x* and *y* are the distances of the breakpoint to respectively the start and end coordinates of the alignment on the reference sequence (see Figure 2), they must satisfy the following condition in equation (1) for the alignment to be kept :

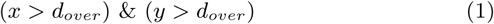

**Figure 2:**
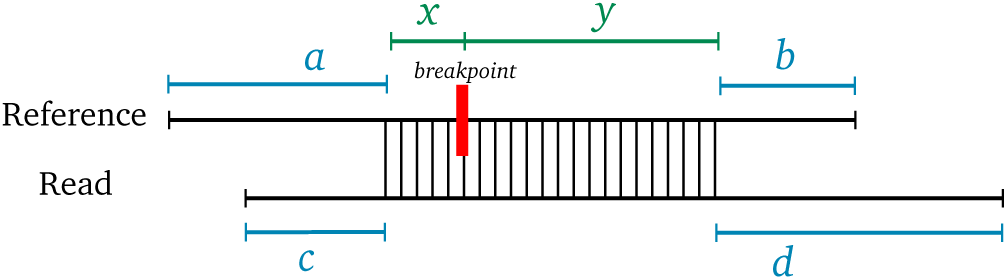
Definition of the different distances on the alignment with respect to the breakpoint (*x* and *y*) and to the sequence extremities (*a, b, c* and *d*) used to select informative alignments. The parts of the sequences that are aligned are illustrated by vertical bars.

Concerning the filtering of spurious false positive alignments, Minimap2 alignments are first filtered based on the quality score. To focus on uniquely mapped reads, the quality score of the alignments must be greater than 10. This is not sufficient to filter out alignments due to repetitive sequences since mapping is performed on a small subset of the reference genome and these alignments may appear as uniquely mapped on this subset.

As Minimap2 is a sensitive local aligner, many of the spurious alignments only cover subsequences of both the reference and the read sequences. To maximize the probability that the aligned read originates from the reference locus, we, therefore, require that the read is aligned to the reference sequence in a semi-global manner. Each alignment extremity must correspond to an extremity of at least one of the two aligned sequences. This criterion gathers four types of situations, namely the read is included in the reference sequence, or *vice-versa*, or the read left end aligns on the reference right end or *vice-versa*.

Indeed this criteria is not strictly applied and a distance of *d*_*end*_ of the alignment to an extremity of at least one of the two aligned sequences is tolerated (*d*_*end*_ is set by default to 100 bp). More formally, if *a* and *b* (resp. *c* and *d*) are the distances of the alignment to the, respectively, left and right extremities of the reference sequence (resp. read sequence) (see Figure 2), then the alignment must fulfill the following condition in equation (2) to be kept:

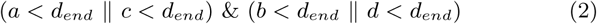

### 2.4 Genotype estimation

For each variant, the genotype is estimated based on the ratio of amounts of reads informatively aligned to each reference allele sequence. Each variant has two reference sequences of different sizes, so even if both alleles are covered with the same read depth, there would be fewer reads that align on the shortest allele sequence. To prevent a bias towards the longest allele, reported read counts for the longest allele are normalized according to the reference allele sequence length ratio, assuming that read count is proportional to the sequence length.

Finally, a genotype is estimated if the variant presence or absence is supported by at least *min_cov* different reads after normalization (sum of the read counts for each allele). By default, this parameter is set to 3.

Genotypes are estimated according to a maximum likelihood strategy. The likelihoods of the three possible genotypes given the observed normalized read counts (*c*_0_ and *c*_1_) are computed based on a simple binomial model, assuming a diploid individual, as described in [15] (see also [16]):

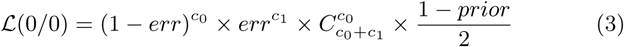

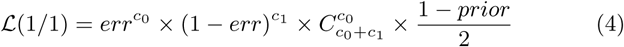

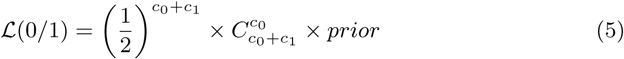

where *err* is the probability that a read maps to a given allele erroneously, assuming it is constant and independent between all observations, and *prior* is the *a priori* probability of the heterozygous genotype. Here, *err* was fixed to 5.10^−5^, after empirical experiments, and the prior distribution is uniform with all three genotypes equally probable 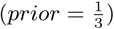. Finally, the genotype with the largest likelihood is assigned and all three likelihoods are also output (-log10 transformed) as additional information in the VCF file.

### 2.5 Implementation and availability

We provide an implementation of this method named SVJedi, freely available at https://github.com/llecompte/SVJedi, under the GNU Affero GPL license. SVJedi is written in Python 3, it requires as input a set of SVs (VCF format), a reference genome (fasta format) and a sequencing read file (fastq or fasta format). Notably, the main steps are implemented in a modular way, allowing the user to start or re-run the program from previous intermediate results. As an example, the first step is not to be repeated if there are several long reads datasets to be genotyped on the same SV set.

## 3 Material

### 3.1 Long read simulated dataset

SVJedi was assessed on simulated datasets on the human chromosome 1 (assembly GRCh37) based on real characterized deletions for the human genome. From the dbVar database [3], we selected 1,000 existing deletions on chromosome 1 (defined as `<DEL>` SV type), which are separated by at least 10,000 bp. The sizes of the deletions vary from 50 bp to 10 kb. In this experiment, deletions were distributed into the three different genotypes: 333 deletions are simulated with 0/0 genotype, 334 deletions with 0/1 genotype and the 333 remaining deletions with 1/1 genotype. Two different sequences were simulated containing each overlapping sets of deletions, representing the two haplotypes of the simulated individual. 1/1 genotype deletions were simulated on both haplotype sequences, whereas deletions of 0/1 genotype were simulated each on one randomly chosen of the two haplotype sequences. Then PacBio data were simulated on both haplotypes, using SimLoRD [17] (version v1.0.2) with varying sequencing error rates (6 %, 10 %, 16 % and 20 %), and at varying total sequencing depths (6x, 10x, 16x, 20x, 30x, 40x, 50x and 60x). Most results presented in the main text are for a 16 % error rate and 30x sequencing depth. Ten such datasets were simulated to assess the reproducibility of results.

### 3.2 Real data

SVJedi was applied on a real human dataset, from the individual HG002, son of the so-called *Ashkenazi trio* dataset. A PacBio sequencing dataset for HG002 was downloaded from the FTP server of GiaB and down-sampled to 30x read depth (FTP links are given in Supplementary Material). We considered the assembly GRCh37.p13 as the human genome reference and as a gold standard call set, we used the SV benchmark set (v0.6) of HG002 individual provided by the GiaB Consortium [18]. This set contains 5,464 high confidence deletions and 7,281 insertions (`PASS` filter tag), whose sizes range from 50 bp to 125 kb (median size of 149 bp and 215 bp for deletions and insertions, respectively).

These SVs were also genotyped in PacBio sequencing datasets of the two parents (HG003 and HG004, 30x and 27x respectively) in order to assess the level of Mendelian inheritance consistency of the son predicted genotypes.

Finally, we considered a real short read dataset for the HG002 individual, 2 × 250 bp Illumina dataset from GiaB, that was down-sampled to 30x read depth. This short read dataset is used for comparison with a short read based SV genotyping approach.

### 3.3 Evaluation

To evaluate the accuracy of the method, a contingency table between the estimated genotypes and the true (simulated) ones is computed, providing a clear view of the number and type of correctly and incorrectly estimated genotypes. The precision of the method is then assessed as the number of correctly estimated genotypes overall all estimated genotypes, as shown in equation (6). The percentage of SVs for which a genotype could be estimated is also measured, and hereafter called the genotyping rate (equation (7)).

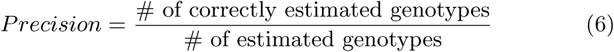

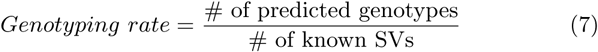

### 3.4 Comparison with other genotyping approaches

We compared our approach with a SV discovery tool that allows SV genotyping, on one of the previously described human chromosome 1 simulated long read dataset. Simulated PacBio reads are first aligned with NGMLR (version 0.2.7) on human chromosome 1. Then, we use Sniffles (version 1.0.11), with default parameters to perform SV discovery and SV genotyping. This analysis does not always predict deletions at the exact simulated coordinates, so a predicted deletion by Sniffles is considered identical as the expected deletion if both deletions overlap by at least 70 %.

Besides, SVJedi was also compared to a SV genotyping approach based on short read data. To do this, the short reads are first aligned with SpeedSeq [4] (version 0.1.2), then the known variants are genotyped with SVtyper (version 0.7.0) with the default settings.

## 4 Results

### 4.1 Assessing SVJedi accuracy on simulated deletions

#### 4.1.1 SVJedi accuracy and robustness

To comprehensively assess the accuracy and robustness of SVJedi, it was first applied to simulated data. Shown results for SVJedi are only for deletion type SVs, as insertions variants are simply the counterpart of deletions. PacBio long reads were simulated on artificial diploid genomes obtained by introducing deletions in the human chromosome 1. Importantly, the sets of introduced and genotyped deletions are made of real characterized deletions in human populations, to reflect the real size distribution and the real complexity of deletion breakpoints and neighboring genomic contexts. To do so, one thousand deletions located on human chromosome 1 were randomly selected from the dbVar database, ranging from 50 to 10,000 bp in size.

Table 1 shows the obtained genotypes compared with expected ones for one simulated dataset at 30x read depth. On this dataset, SVJedi achieves 97.8 % precision, with 974 deletions correctly predicted over 996 with an assigned genotype. Among the 1,000 assessed deletions, only 4 could not be assigned a genotype due to insufficient coverage of informative reads, the genotyping rate being thus 99.6 %. Among the few genotyping errors, most concern 1/1 genotypes that were incorrectly predicted as 0/1.

**Table 1:** Contingency table of SVJedi results on PacBio simulated data (30x) of human chromosome 1 with 1,000 deletions from dbVar. SVJedi genotype predictions are indicated by column and the expected genotypes are shown by row. The genotype “./.” column corresponds to deletions for which SVJedi could not assess the genotype.

|  |  | SVJedi predictions |  |  |  |
| --- | --- | --- | --- | --- | --- |
|  |  | 0/0 | 0/1 | 1/1 | ./. |
| Truth | 0/0 | 331 | 1 | 0 | 1 |
|  | 0/1 | 3 | 330 | 0 | 1 |
|  | 1/1 | 0 | 18 | 313 | 2 |

Precision : 97.8 %

SVJedi precision results were evaluated in terms of varying sequencing depths, ranging from 6x to 60x (see Fig. 3). As expected, the accuracy of SVJedi increases with the read depth, but interestingly, even at low coverage (6x) the accuracy is on average above 94 % and a plateau is quickly reached between 20x and 30x, with already 97 % of precision at 20x.

**Figure 3:**
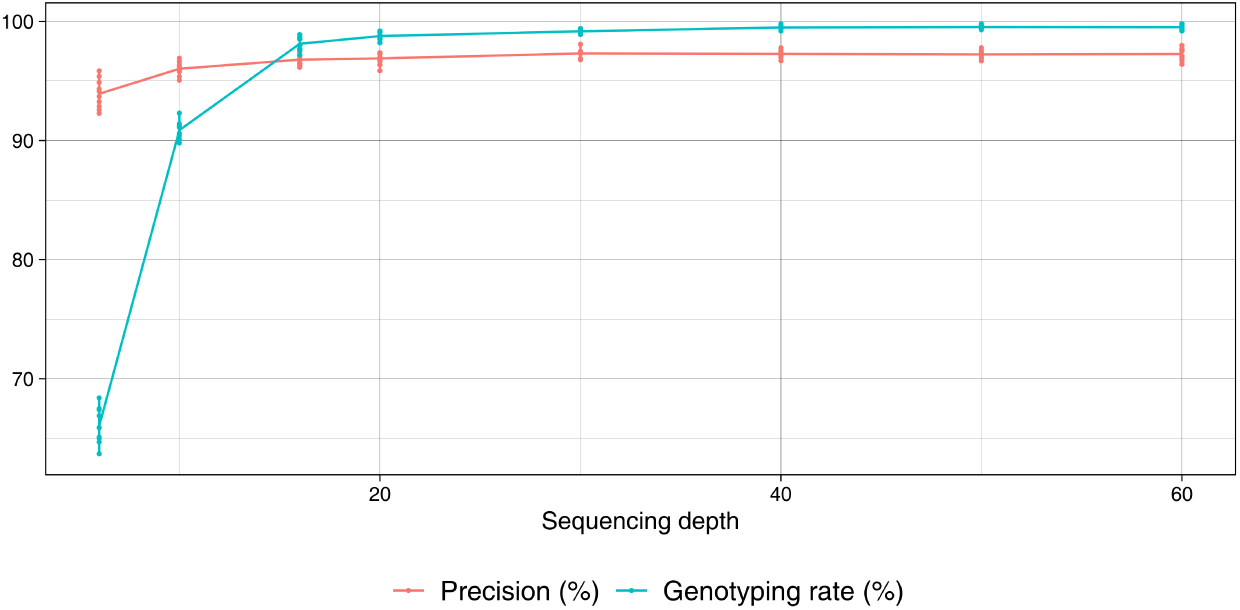
SVJedi precision results as a function of the sequencing depth for ten simulated PacBio datasets of human chromosome 1, containing 1,000 deletions from the dbVar database. The red dots correspond to the average precision and the red segments represent the standard deviations, at each sequencing depth.

Similarly, SVJedi results were evaluated in terms of varying sequencing error rates. In this case, both precision and genotyping rate were not impacted by a lower or higher sequencing error rate as long as it stays realistic (see Supplementary Figure 1).

The breakpoint coordinates of SVs detected by SV discovery methods are not always defined at the base pair resolution. In order to assess to what extent this potential imprecision can impact the genotyping precision of SVJedi, we performed experiments with altered breakpoint positions in the input variant VCF file. All breakpoint positions have been randomly shifted according to a Normal distribution centered on the exact break-point position with several standard deviations (*σ*) values ranging from 10 to 100 bp. We show that the genotyping accuracy with *σ* equals 50 bp does not fall below 94 % (see Supplementary Figure 2), indicating that SVJedi is not much impacted by the exact definition of the positions of the reference breakpoints.

#### 4.1.2 Comparison with a SV discovery approach

One can wonder if these simulated deletions could be easily detected and genotyped by a long read SV discovery tool. We applied here one of the best to date such tool, Sniffles [12, 19], to the chromosome 1 simulated read dataset. As expected, none of the 333 simulated deletions with 0/0 genotypes were assigned a genotype in the Sniffles output call set, since a discovery tool naturally only reports actual differences with the reference genome. As a result, among the 667 deletions simulated with either a 0/1 or 1/1 genotype, only 570 were discovered by Sniffles, giving a recall of 85.5 %, with mainly the heterozygous genotypes that are missing (24 % of 0/1 deletions were missed, versus 5 % for the homozygous ones). Interestingly, Sniffles often badly estimates the genotype of the discovered deletions, assigning most of the 1/1 discovered deletions (n = 247, 74 %) as heterozygous, and resulting in a genotyping precision of only 53.3 %. Detailed results are given in Supplementary Table 1. This highlights the fact that Sniffles, a SV discovery tool, is much less precise for the genotyping task than a dedicated genotyping tool.

### 4.2 Application to a real human dataset

#### 4.2.1 SVJedi results on HG002 individual

To get closer to the reality of biological data, we applied our tool to a real human dataset, the HG002 individual, son of the so-called *Ashkenazi trio* dataset, which has been highly sequenced and analyzed in various benchmarks and notably by the GiaB Consortium [20]. The latter, precisely, provides a set of high confidence SV calls together with their genotype in the individual HG002. SV discovery and genotyping were based on several sequencing technologies, SV callers and careful call set merging. Their work estimated genotypes for 5,464 deletions and 7,281 insertions, which can then be considered as the ground truth. It should be noted that we can focus only on heterozygous (0/1) and homozygous for the alternative allele (1/1) genotypes. Indeed, the SV call set was obtained from SV discovery methods, which can only detect variations between the individual and the reference genome. SVJedi was applied on a 30x PacBio long read dataset from individual HG002, to assess the genotypes of both deletions and insertions of this high confidence set.

SVJedi results are indicated in the Table 2 for deletions and insertions in the left and right tables, respectively. We observe a good overlap of the predicted genotypes between our genotyping tool and the GiaB call set, illustrated by the grey labeled boxes. More precisely, among the assigned genotypes, there are 91.9 % of deletions and 92.8 % of insertions that are genotyped by SVJedi identically as the GiaB call set.

**Table 2:**
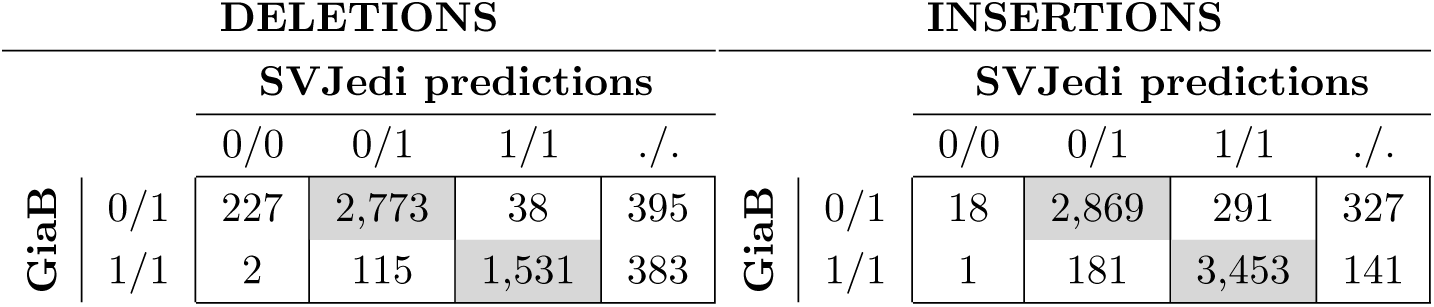
Contingency tables of SVJedi genotyping results on the real 30x PacBio dataset of human individual HG002 with respect to the high confidence GiaB call set. Results for the 5,464 deletions (left) and 7,281 insertions (right) are indicated in two separated tables, where columns indicate SVJedi genotypes and rows GiaB ones. Grey labelled boxes correspond to identical predictions between the two methods. The number of genotypes that SVJedi fail to assess is indicated by the “./.” column.

Compared to previous results on simulated data, SVJedi shows a lower genotyping rate on this real dataset, for both deletions and insertions (85.8 % and 93.6 %, respectively). We notice that the majority of the not genotyped variants are mostly small variants. Indeed, 87.1 % are of size less than 100 bp. The latter seems to be more impacted by the heterogeneity of PacBio sequencing depth since when using the full 63x dataset, the genotyping rate increases to 96.9 % (94.3 % and 98.4 % for deletions and insertions, respectively).

Similarly, among the SVs differently genotyped between SVJedi and GiaB, a large part is represented by small variants (56.6 % are smaller than 100 bp). In these cases of small variants, when they are assumed to be heterozygous in the GiaB call set, we can observe that SVJedi tends to favor the largest allele (1/1 for insertions and 0/0 for deletions). This may be explained by the fact that most sequencing errors of PacBio sequencing technology are insertions, that can be wrongly mapped to the largest allele when the latter is small.

Since sequencing data are available for the parents of the studied individual (HG003 for the father and HG004 for the mother), we can check, as an alternative validation approach, if the predicted genotypes for the son are consistent with his parent genotypes, assuming perfect Mendelian inheritance and a very low de novo mutation rate.

To do so, from the same set of deletions and insertions, which is the GiaB call set, SVJedi was applied to three PacBio sequence datasets, one per individual, with a sequencing depth of about 30x for each dataset. Overall, the Mendelian inheritance consistency of SVJedi on this trio dataset is high, with 96.9 % of the son genotypes that are consistent with his parent genotypes. As expected, most inconsistent genotypes concern SVs that were genotyped differently between SVJedi and GiaB (48.7 %, n = 154), confirming for those that they are probably wrongly assessed by SVJedi. However, these confirmed errors represent only 1.2 % of the dataset.

#### 4.2.2 Comparison with a short read based genotyping approach

For this same individual (HG002), some short read datasets are also available, we, therefore, can compare SV genotyping performances between two approaches and data types, namely long versus short reads. SVJedi predictions were compared to a SV genotyping tool for short reads, SVtyper, known as a reference tool in the state of the art [4, 7]. Since SVtyper does not support insertion variants, we focus here only on deletions, and the 5,464 deletions from the GiaB call set were genotyped with SVtyper using a 2×250 bp 30x Illumina read dataset of HG002.

Table 3 shows that more than half of the deletions are genotyped differently by SVtyper than in the high confidence GiaB call set, resulting in a genotyping precision of 46.4 %, while this percentage rises to 92.4 % for SVJedi with long reads. Remarkably, many of the discrepancies of SVtyper with GiaB are totally contradictory with 0/0 genotypes instead of 1/1 ones. This demonstrates the higher benefit of using long reads and a dedicated genotyping tool such as SVJedi rather than short reads.

**Table 3:** SVtyper genotyping results on a real 30x Illumina dataset of HG002 in columns compared to the high confidence GiaB genotype calls in rows. Grey labelled boxes correspond to identical predictions between the two methods. Last column, “./.”, corresponds to deletions for which SVtyper failed to assess the genotype.

|  |  | SVtyper predictions |  |  |  |
| --- | --- | --- | --- | --- | --- |
|  |  | 0/0 | 0/1 | 1/1 | ./. |
| GiaB | 0/1 | 1862 | 1550 | 13 | 8 |
|  | 1/1 | 803 | 230 | 962 | 36 |

Importantly, SVJedi does not come with a high computational cost. Genotyping the 12,745 SVs with the 30x PacBio HG002 dataset took only 2h30m on a Linux 40-CPU node running at 2.60 GHz. The alignment step is the most time-consuming step and took 2h20m. These running times are comparable with short read genotypers, for instance, SVtyper (4h29m), for which similarly most of the time is monopolized by the mandatory mapping step.

## 5 Discussion and conclusion

In conclusion, we provide a novel SV genotyping approach for long read data, that showed good results on simulated and real datasets for both deletions and insertions. The robustness of our tool, SVJedi, was also highlighted in this work, for several sequencing depths and error rates, but also to the precision of the breakpoint positions. The approach is implemented in the SVJedi software for the moment for the most common and studied types of SVs, deletion and insertion variants, which represent to date 99 % of dbVar referenced SVs. This proof of principle is the first step before generalizing the approach for all other types of SVs.

This work also demonstrated that this is crucial to develop dedicated SV genotyping methods, as well as SV discovery methods. Firstly, because this is the only way to get evidence for the absence of SVs in a given individual. Secondly, and more surprisingly, because SV discovery tools are not as efficient and precise to genotype variants once they have been discovered, at least with long read data as was shown here. Indeed, without a priori SV discovery is a much harder task than genotyping SVs with well-characterized alleles, but when the aim strictly is to genotype or compare individuals on already known variants, we have shown that using as much as possible the known features of variants is much more efficient.

Also, on real human data, we were able to quantify the impact of the sequencing technology on SV genotyping. Although this was expected that long read data would perform better than short read ones, the observed difference is considerable with a twofold increase of the precision with long reads. This is in particular due to the very poor performances obtained with short reads, that are ill-adapted to deal with the complex and repeat-rich regions often present at SV junctions. On the opposite, this work shows that the long-distance information contained in long reads can be efficiently used to discriminate between breakpoints, despite relatively high sequencing error rates and variability in sequencing coverage. This result underlines the relevance of such a method dedicated to genotyping from long read data.

Although long read sequencing technology remains to date more expensive than short read ones, to be used for instance in routine in the clinical setting [21], we can hope that this situation will improve in the next few years. The high precision and low computational requirements of SVJedi make it ready for such happening and to be integrated into routine pipelines to screen for instance disease-related SVs and therefore improve medical diagnosis or disease understanding.

## Supporting information

SVJedi_supplementary

## Acknowledgements

We are thankful to the Genouest bioinformatics platform, computations have been made possible thanks to the resources of the Genouest infrastructure.

## References

[1] Peter A Audano, Arvis Sulovari, et al. Characterizing the major structural variant alleles of the human genome. Cell, 176(3):663–675, 2019.

[2] James R Lupski. Structural variation mutagenesis of the human genome: Impact on disease and evolution. Environ. Mol. Mutagen., 56(5):419–436, 2015.

[3] Lon Phan, Jeffrey Hsu, et al. dbVar structural variant cluster set for data analysis and variant comparison. F1000Research, 5, 2017.

[4] Colby Chiang, Ryan M Layer, et al. SpeedSeq: ultra-fast personal genome analysis and interpretation. Nat. Methods, 12(10):966–968, 2015.

[5] Danny Antaki, William M Brandler, et al. SV2: accurate structural variation genotyping and de novo mutation detection from whole genomes. Bioinformatics, 34(10):1774–1777, 2017.

[6] Parsoa Khorsand and Fereydoun Hormozdiari. Nebula: Ultra-efficient mapping-free structural variant genotyper. bioRxiv, 2019.

[7] Varuna Chander, Richard A Gibbs, et al. Evaluation of computational genotyping of structural variation for clinical diagnoses. GigaScience, 8(9), 2019.

[8] Alexis L Norris, Rachael E Workman, et al. Nanopore sequencing detects structural variants in cancer. Cancer Biol. Ther., 17(3):246–253, 2016.

[9] John Huddleston, Mark JP Chaisson, et al. Discovery and genotyping of structural variation from long-read haploid genome sequence data. Genome Res., 27(5):677–685, 2017.

[10] M Cretu Stancu, Markus J Van Roosmalen, et al. Mapping and phasing of structural variation in patient genomes using nanopore sequencing. Nat. Commun., 8(1):1326, 2017.

[11] Miten Jain, Sergey Koren, et al. Nanopore sequencing and assembly of a human genome with ultra-long reads. Nat. Biotechnol., 36(4):338, 2018.

[12] Fritz J Sedlazeck, Philipp Rescheneder, et al. Accurate detection of complex structural variations using single-molecule sequencing. Nat. Methods, 15(6):461–468, 2018.

[13] David Heller and Martin Vingron. SVIM: structural variant identification using mapped long reads. Bioinformatics, 35(17):2907–2915, 2019.

[14] Heng Li. Minimap2: pairwise alignment for nucleotide sequences. Bioinformatics, 34(18):3094–3100, 2018.

[15] Rasmus Nielsen, Joshua S Paul, et al. Genotype and SNP calling from next-generation sequencing data. Nat. Rev. Genet., 12(6):443, 2011.

[16] Heng Li. A statistical framework for SNP calling, mutation discovery, association mapping and population genetical parameter estimation from sequencing data. Bioinformatics, 27(21):2987–2993, 2011.

[17] Bianca K Stöcker, Johannes Köster, et al. SimLoRD: Simulation of Long Read Data. Bioinformatics, 32(17):2704–2706, 2016.

[18] Justin M Zook, Nancy F Hansen, et al. A robust benchmark for germline structural variant detection. bioRxiv, 2019.

[19] Wouter De Coster, Peter De Rijk, et al. Structural variants identified by Oxford Nanopore PromethION sequencing of the human genome. Genome Res., 29(7):1178–1187, 2019.

[20] Justin M Zook, David Catoe, et al. Extensive sequencing of seven human genomes to characterize benchmark reference materials. Sci. Data, 3:160025, 2016.

[21] Jason D Merker, Aaron M Wenger, et al. Long-read genome sequencing identifies causal structural variation in a Mendelian disease. Genet. Med., 20(1):159, 2018.

