## Supplementary material for "SVJedi: Genotyping structural variations with long reads": SVJedi_supplementary

### SVJedi: Genotyping structural variations with long reads
#### Supplementary material

Lolita Lecompte, Pierre Peterlongo, Dominique Lavenier and Claire Lemaitre

#### 1 Accessibility to real data

**The gold standard call set** provided by Genome in a Bottle (GiaB) Consortium is available at the following link:

- `ftp://ftp-trace.ncbi.nlm.nih.gov/giab/ftp/data/AshkenazimTrio/analysis/NIST_SVs_Integration_v0.6/HG002_SVs_Tier1_v0.6.vcf.gz`

In our study, we focus only on variants of the Tier 1 call set which are isolated and sequence-resolved SVs, corresponding to the `PASS` filter of this VCF file. This call set represents 5,464 deletions and 7,281 insertions.

**Real Pacific Biosciences (PacBio) sequence datasets** for Ashkenazi trio individuals provided by GiaB are available at the following links:

- `ftp://ftp-trace.ncbi.nlm.nih.gov/giab/ftp/data/AshkenazimTrio/HG002_NA24385_son/PacBio_MtSinai_NIST/`
- `ftp://ftp-trace.ncbi.nlm.nih.gov/giab/ftp/data/AshkenazimTrio/HG003_NA24149_father/PacBio_MtSinai_NIST/`
- `ftp://ftp-trace.ncbi.nlm.nih.gov/giab/ftp/data/AshkenazimTrio/HG004_NA24143_mother/PacBio_MtSinai_NIST/`

The PacBio data of the individual HG002 has a sequencing depth of 63x. The sequence data were sub-sampled using SAMtools [1] to a depth of 30x.

**Real Illumina dataset for HG002** is available at the following link:

- `ftp://ftp-trace.ncbi.nlm.nih.gov/giab/ftp/data/AshkenazimTrio/HG002_NA24385_son/NIST_Illumina_2x250bps/`

As with the PacBio data sample of the individual HG002, the Illumina data sample was sub-sampled to a sequencing depth of 30x as well. The data was then aligned with SpeedSeq on the human reference genome GRCh37.p13 and genotyped with SVtyper with default parameters. Both `CIPOS` and `CIEND` were set to 0,0 for all variants in the VCF before running SVtyper.

```
speedseq align -t 40 -M 20 \
    -R "@RG\tID:HG002\tSM:Illumina\tLB:2x250" \
    -o speedseq_aln_HG002_Illumina \
    GRCh37_latest_genomic.fna reads.end1.fq reads.end2.fq
```

**Simulated PacBio reads with SimLoRD** Long reads are simulated on two distinct haplotypes where the deletions from dbVar are added according to the simulated genotype (0/0, 0/1, 1/1). The SimLoRD commands are shown below for a dataset with a sequencing depth of 30x and a sequencing error rate of 16 %. The same distribution of errors between insertions, deletions and substitutions, as LRCstats [2] has been defined.

```
simlord --read-reference haploA_with_1000_simulated_DEL.fasta --coverage 15
    --max-passes 1 -pi 0.11 -pd 0.04 -ps 0.01 simulated_reads_haploA

simlord --read-reference haploB_with_1000_simulated_DEL.fasta --coverage 15
    --max-passes 1 -pi 0.11 -pd 0.04 -ps 0.01 simulated_reads_haploB
```

### 2 Assessment of the SVJedi's robustness

#### 2.1 SVJedi results on simulated PacBio dataset for several error rate

The robustness of SVJedi is evaluated on simulated data at 30x sequencing depth based on several error rates: 6 %, 10 %, 16 %, 20 %.

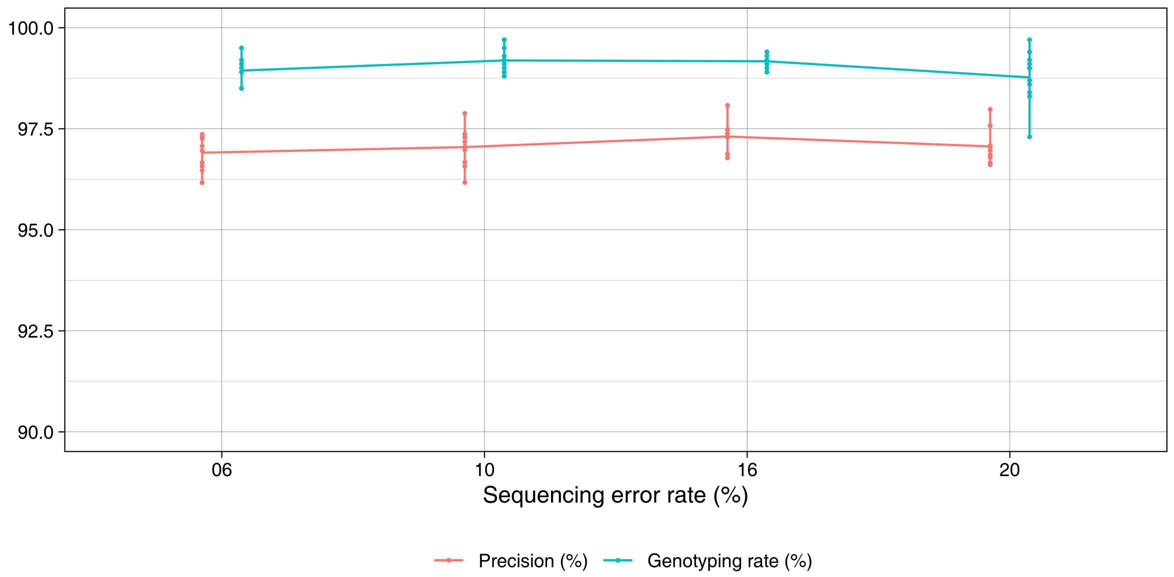


Supplementary Figure 1: SVJedi precision results on simulated 30x datasets at multiple sequencing error rate : 6 %, 10 %, 16 %, 20 %).

#### 2.2 SVJedi results on simulated PacBio dataset with shifted breakpoints

The robustness of SVJedi is evaluated on simulated data at 30x sequencing depth according to the definition of breakpoint positions. Breakpoint positions are altered in the input variant VCF file. All breakpoint positions have been randomly shifted according to a Normal distribution centered on the exact breakpoint position with several standard deviations values ranging from 10 to 100 bp.

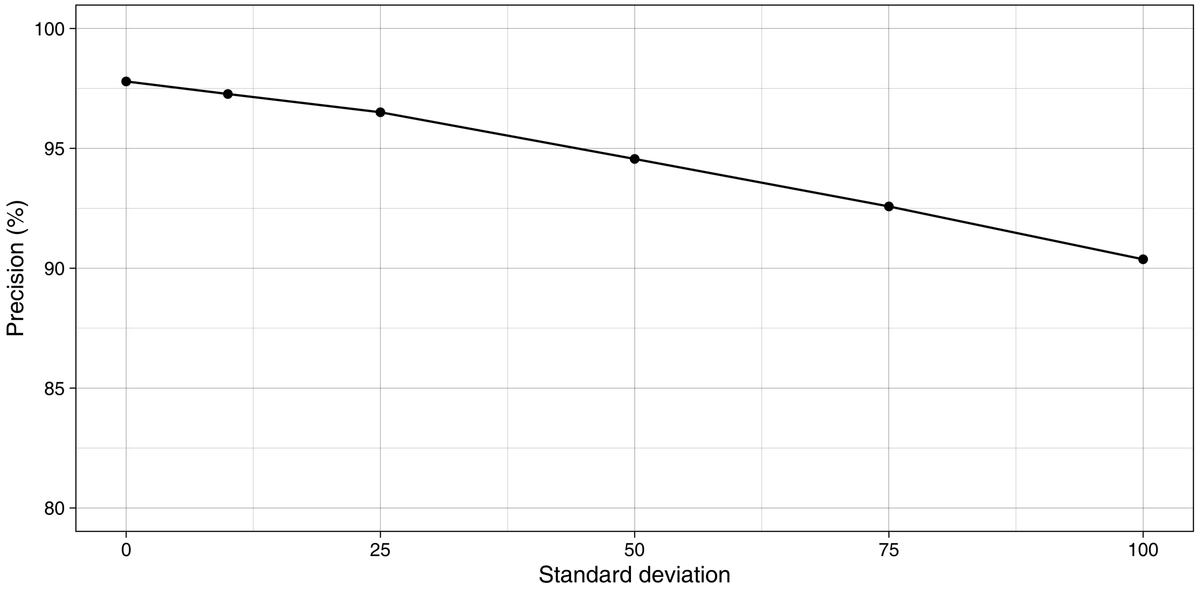


Supplementary Figure 2: SVJedi precision results on simulated 30x dataset at multiple shifted breakpoints position according to a Nomal law, with several standard deviations values ranging from 10 bp to 100 bp

Supplementary Figure 2 shows the SVJedi precision as a function of the offset of breakpoint positions for several standard deviations. It can be seen that at a standard deviation of 50, the SVJedi accuracy is greater than 94 %.

### 3 Sniffles prediction results on simulated dataset

Sniffles command SV discovering and genotyping:

```
sniffles -m 30x_PacBio_mapped.sorted.bam -v output.vcf --genotype
```

| | | Sniffles predictions | | | |
|---|---|---|---|---|---|
| | | 0/0 | 0/1 | 1/1 | ./. |
| Truth | 0/1 | 19 | 234 | 0 | 81 |
| | 1/1 | 0 | 247 | 70 | 16 |

Precision = 53.3 %

Supplementary Table 1: Contingency table of Sniffles results on PacBio simulated data (30x). Sniffles genotype predictions are indicated by column and the expected genotypes are shown by row. The genotype "./." column corresponds to deletions for which Sniffles was not able either to detect the deletion or to assess its genotype.

Sniffles being a tool for detecting SV, homozygous SVs for the reference allele (0/0) cannot be detected or genotyped. Thus, in Supplementary Table 1, we are only interested in heterozygous (0/1) or homozygous SVs for the alternative allele (1/1).

#### References


[1] Heng Li, Bob Handsaker, et al. The sequence alignment/map format and SAMtools. *Bioinformatics*, 25(16):2078–2079, 2009.

[2] Sean La, Ehsan Haghshenas, et al. LRCstats, a tool for evaluating long reads correction methods. *Bioinformatics*, 33(22):3652–3654, 2017.
